# Role of bacterial persistence in spatial population expansion

**DOI:** 10.1101/2021.03.16.435668

**Authors:** Pintu Patra, Stefan Klumpp

## Abstract

Bacterial persistence, tolerance to antibiotics via stochastic phenotype switching provides a survival strategy and a fitness advantage in temporally fluctuating environments. Here we study its possible benefit in spatially varying environments using a Fisher wave approach. We study the spatial expansion of a population with stochastic switching between two phenotypes in spatially homogeneous conditions and in the presence of an antibiotic barrier. Our analytical results show that the expansion speed in growth-supporting conditions depends on the fraction of persister cells at the leading edge of the population wave. The leading edge contains a small fraction of persister cells, keeping the effect on the expansion speed minimal. The fraction of persisters increases gradually in the interior of the wave. This persister pool benefits the population when it is stalled by an antibiotic environment. In that case, the presence of persister enables the population to spread deeper into the antibiotic region and to cross an antibiotic region more rapidly. The interplay of population dynamics at the interface separating the two environments and phenotype switching in the antibiotic region results in a optimal switching rate. Overall, our results show that stochastic switching can promote population expansion in the presence of antibiotic barriers or other stressful environments.

## INTRODUCTION

Population expansion in space facilitates evolutionary diversification and survival of species [1, 2]. Recent experiments using microfluidics have demonstrated the role of spatial structures on population expansion using microbes as model organisms, providing insight into several eco-evolutionary [3–6] and medically relevant questions [7–10]. For example, in spatial environments, cooperative behaviors are sustained [5], positively frequencydependent selection can persist [11], and the rapid emergence of antibiotic resistance is facilitated [12]. Population survival and competitive strategies are the major driving factors for many of these intriguing behaviors [5, 9, 13].

A prime example of a population-level survival strategy is bacterial persistence, where the population benefits from a subpopulation with the persister phenotype that is more tolerant to stresses such as antibiotics [14–16]. However, persisters incur a cost due to their slow division rate in growth-supporting conditions [17]. In a temporally changing environment, the interplay between cost and benefit determines the circumstances where persistence is advantageous [18–21]. A population expanding in space could encounter such temporally varying conditions by moving through different environments in space. Therefore we ask here how bacterial persistence affects the expansion of a population in space and whether a similar cost-benefit trade-off exists in spatially modulated environments as in temporally varying conditions.

To this end, we study the effect of bacterial persistence in a population spatially expanding in a homogeneous growth-favorable environment and in a scenario where growth is halted by an antibiotic region. We make use of the approach introduced by Fisher [22] and Skellam [23] which has been extensively used to describe the spatial spread of invading species, insects, epidemic agents, and so on [24]. This formalism allows us to write a set of wave equations for a population of cells that reversibly switch between the normal growing state and the slow-growing persister state. We determine the cost of persistence during growth conditions by computing the population expansion speed as well as the fitness advantage due to persisters in crossing an antibiotic barrier. The scenarios studied here can be considered idealized descriptions of real environments, for example in the body of a patient, but more importantly they can directly be realized experimentally with current methods such as spatially structured environments on solid surfaces [7] or in microfluidic devices [9].

## MODEL FOR SPATIAL EXPANSION OF A PHENOTYPICALLY HETEROGENEOUS POPULATION

The expansion of populations in space can be described by the Fisher equation (or Kolmogorov–Petrovsky–Piskunov equation)

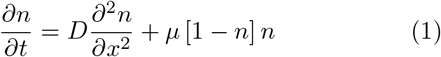

Here *n*(*x, t*) is the density of a population at position *x* at time *t*. To keep the model simple, we consider spatial expansion in one dimension. The two terms on the right hand side of the equation describe the diffusive spread of the population in space (with a mobility or diffusion coefficient *D)* and its local logistic growth with growth rate *μ*. Note that the population size is normalized to the carrying capacity of the logistic growth, i.e. to the maximum population size that can be achieved in the spatially homogeneous conditions. In the following we extend this equation to the case of a population with two distinct phenotypes using bacterial persistence as a concrete example. Bacterial persistence in the presence of antibiotics is associated with an intrinsic phenotypic heterogeneity in the bacterial population. This heterogeneity arises from the stochastic transition between the normal growing cell state and the drug-tolerant persister state at the single-cell level [17]. The two phenotypic states are characterized by different growth (in growth conditions) and death rates (in the presence of antibiotics). To include such phenotype switching into the model, we describe the normal cells and the persister cells by densities *n*(*x,t*) and *p*(*x,t*), respectively. They are characterized by different growth rates *μ_n_* and *μ_p_* and are subject to a common carrying capacity, such that the constraint on the population size due to the logistic growth acts on the sum *n* + *p*. In addition, we allow for different mobility parameters *D_n_* and *D_p_*. A difference in mobility can be expected if movement is due to self-propulsion; if, however, movement is driven by external driving forces, the two parameters will likely be the same. Finally we include phenotype switching: A cell in the normal state can switch to the persister state with a rate *a* and vice-versa with a rate *b.* These rates are typically small compared to the growth rate [17].

Taking these considerations together, the dynamics of the population composed of normal cells (*n*) and persister cells (*p*) can be described by the following coupled differential equations,

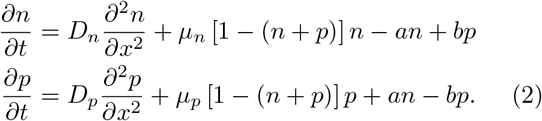

Like the well-known Fisher equation, these equations display a traveling wave solution. The numerical solution of the above equations is shown in Figure 1 and exhibits a traveling wave for both subpopulations. The numerical solution shows that the fraction of persister cells is small at the leading edge and increases progressively towards the interior of the wave. Far from the leading edge, the subpopulation sizes are determined by phenotype switching alone and given by *p* ≈ *a*/(*a* + *b*) and *n* ≈ *b*/(*a* + *b*) (in Figure 1, we use equal switching rates (*a* = *b*), which results in equal subpopulation densities, *n = p* = 0.5, in the interior of the wave). Next, we compute the speed of the population wave as a function of the switching rates. We find that the wave speed decreases with increasing switching rate, while concurrently the persister fraction at the leading edge of the wave increases as shown in Figure 1b.

**FIG. 1:**
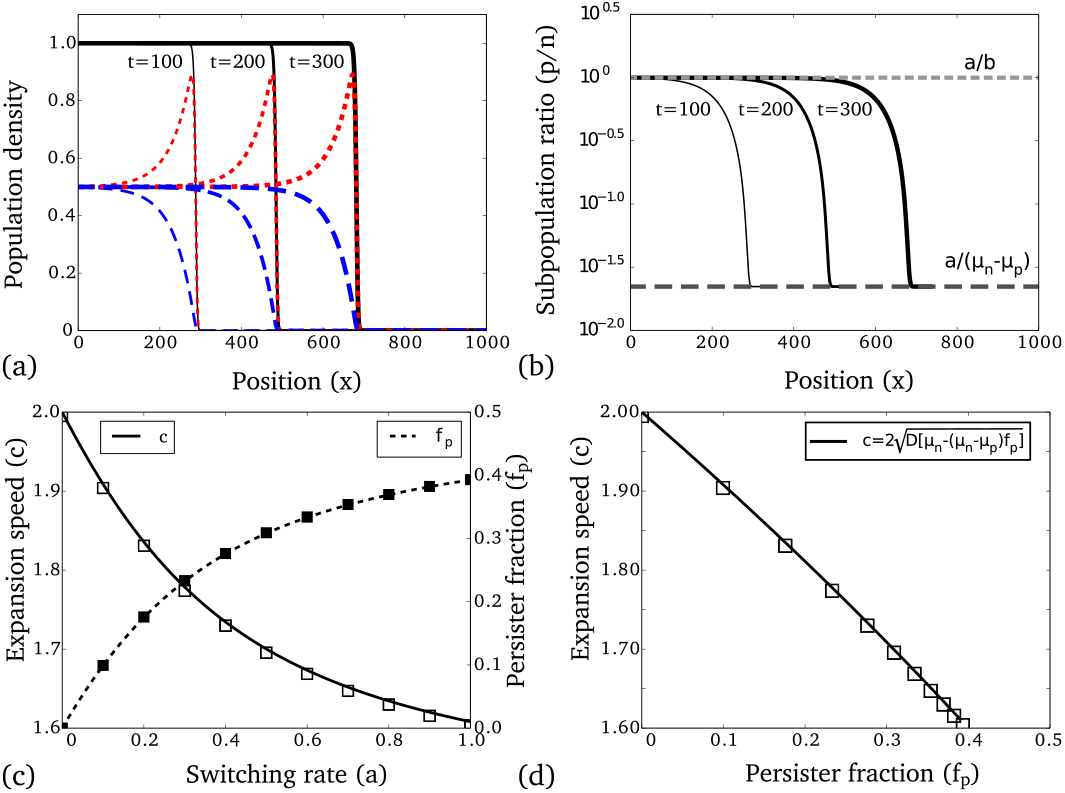
(a) The model equations display traveling waves moving with a constant expansion speed. The total population density (black line) wave has contributions from two subpopulation waves corresponding to normal (red broken line) and persister cells (blue dashed line). We have used the following parameters for numerical calculations: *μ_n_* = 1, *μ_n_* = 0.1, *D* = 1, *b* = *a* = 0.02. (b) The subpopulation ratio (*p/n*) at the leading edge and back of the wave follows the theoretically predicted ratios for the exponential growth (dark grey dashed line) and stationary phase (grey dashed line), respectively. (c) Expansion speed (open squares) and persister fraction at the leading edge (closed squares) as functions of the phenotype switching rates (for *a = b*). The solid lines show the corresponding analytical results. (d) The decrease in the expansion speed with the increase in persister fraction at the leading edge (open squares) follows the constitutive relation given by equation 4.

## RESULTS

### Characteristic features of the population front

The characteristic features of the population front as presented in Figure 1 can be understood based on the subpopulation balance during exponential growth and stationary phase in non-moving conditions. In the absence of movement (diffusion rates are *D_n_* = *D_p_* = 0), during exponential growth phase when the population size is below the carrying capacity (where *n + p* ≪ 1), the ratio of persisters to the normal subpopulation is approximately given by *p/n* ≈ *a*/(*μ_n_* — *μ_p_*) [20]. The latter expression follows from a balance of two effects: normal cell outgrow persisters with rate *μ_n_* — *μ_p_*, but also regenerate them through switching. In the stationary phase when the population size is near the carrying capacity (*n + p* ≈ 1), the ratio of persisters to the normal subpopulation is *p/n* ≈ *a/b* [20]. In a spatially expanding population (i.e., for finite diffusion rate), the front of the population exhibits the exponential scenario, while the population is in stationary phase far behind the population front (shown in Figure 1b).

For the classical Fisher wave, the expansion speed is determined by the growth and diffusion via the relation 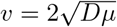. In our case, the growth rate is modulated by the presence of persister cells. This can be demonstrated by deriving an equation for the total population density (*P* = *n + p*) as

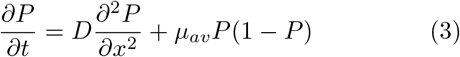

where *μ_av_ μ_n_* (*μ_n_ μ_p_*)*f_p_* and *f_p_* = *p*/ (*n* + *p*) is the fraction of persisters. Hence, in the presence of phenotype switching, the expansion speed is modulated due to a finite fraction of persisters at the tip of the population wave. We validated this numerically by measuring the wave speed and persister fraction simultaneously at the tip of the wave (as shown in Figure 1c). The wave speed (denoted by c) and persister fraction (*f_p_*) at the wave front follow the relation (shown in Figure 1d)

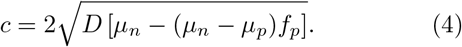

For small phenotype switching rates, the subpopulation ratio at leading edge can be approximated by that for the exponential growth phase, *f_p_* ≈ *a*/(*μ_n_* – *μ_p_*) and the wave speed is given by 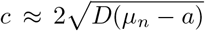. In the limit (*a* → 0), one recovers the maximal expansion speed (corresponding to a single population without persisters). Thus, since typically switching rates are small compared to the growth rate, the effect of persisters on the expansion speed is very small, even if a large subpopulation of persisters exists in the stationary phase situation far from the expansion front.

### Exact expansion speed

Next, we use standard traveling wave analysis to obtain an analytical expression for the expansion speed over a large range of parameters (for the complete analysis see supporting information). This method is based on the observation that at long times these traveling waves propagate with constant speed and fixed front shape. This reduces the partial differential equations into an ordinary differential equation with a single variable *z* = (*x — ct*), the moving frame of reference. This approach has been used to estimate the expansion speed in reaction-diffusion equations describing population expansion, e.g., in bacterial colony expansion models [25, 26], in the dynamics of horizontally transmitted traits [27] and cooperative alleles [28] in expanding population waves. Specifically, we use the following ansatz for the subpopulation densities, *n*(*x,t*) = *n*(*x — ct*) and *p*(*x,t*) = *p*(*x — ct*) in Equation (1) and use the resulting equations to determine the stability of the fixed point (*n* = 0, *p* = 0). The eigenvalues around the fixed point provide a condition for the existence of non-negative and non-oscillatory solutions that determines the minimal value for the wave speed c. Our analysis revealed that traveling waves exist when the following condition (from the requirement that eigenvalues must be real) is satisfied,

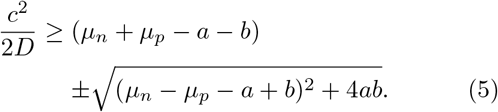

For equal rates of phenotype switching (*a = b*), the condition for the expansion speed simplifies to 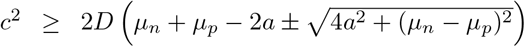. The above condition leads to two distinct values for the expansion speed for which a traveling wave solution exists. We call these the slow and fast mode. The slow mode corresponds to a population that contains mostly persister cells, while the fast mode corresponds to a population consisting mostly of normal cells. This can been seen in the limit *a* = *b* = 0, when the two subpopulation waves decouple with marginal speed 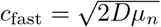 and 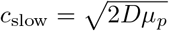. In the coupled case (i.e. with finite switching), the expansion speed is given by the faster one of the two moving fronts, i.e. the positive root of the above expression. However, in some cases, the slow mode will also be relevant. An example is a population after crossing an antibiotic zone. Such a population consists almost exclusively of persisters. It will therefore first ex-pand as a slow-mode wave, which will eventually be taken over by a faster wave of normal cells (generated from persister cells that switch to the normal phenotype). This behavior is shown in Figure 2(a) where the simulation is started with a finite number of persister cells and no normal cells. As the population size increases, the fraction of normal cells increases (shown in Figure 2(b)), and the whole population advances with the fast mode.

**FIG. 2:**
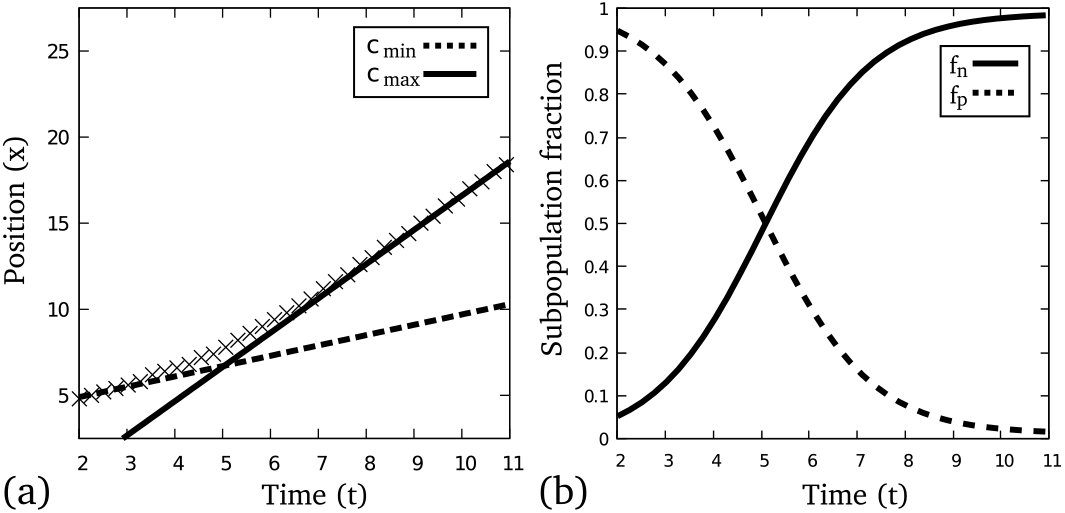
(a) Expansion speed transitions from a slow (persister-dominated) to a fast (normal cell-dominated) mode as seen in our numerical simulations (cross points) that start with a high persister fraction. The front position as a function of time exhibits the two linear regimes with slopes that match the analytic expressions for the fast (solid line) and slow (broken line) expansion speed. (b) During the speed transition, the fractions of the two phenotypes change; the population shifts from a persister-dominated (broken line) to a normal-cell dominated (solid line) wave.

Comparison of Equation (5) with Equation (4) also provides an expression for the persister fraction at the leading edge:

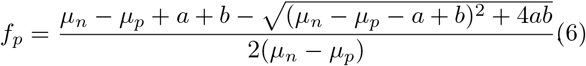

The expressions for the expansion speed c and the persister fraction *f_p_* show excellent agreement with the respective numerically calculated values (shown in Figure 1c). The persister fraction *f_p_* reduces to *a*/(*μ_n_* — *μ_p_*) for small switching rates (where *ab* ≈ 0) in agreement with the fractions expected in exponential phase for spatially homogeneous populations [20].

### Extent of penetration of the heterogeneous wave into an antibiotic region

The above results show that spatial expansion results in a small fraction of persisters at the tip of the wave and a larger fraction in the wave interior. This serves as an advantageous trait in the absence of stresses such as antibiotics, as the population’s expansion is not negatively impacted by persisters, while a persister pool is build up behind the expanding front. If such an expanding population encounters a region that is detrimental to growth, e.g. because of the presence of an antibiotic, the expansion will stall. Population stalling at the interface of an unfavorable environment is known to play a crucial role in the emergence of antibiotic resistance [7], where a key determinant is population survival inside antibiotic environment [29]. Under such stalling conditions, the persister fraction will slowly increase and cells from the stalled population will enter the unfavorable environment, where the population density decays.

To quantify the benefit of bacterial persistence in such scenario, we consider a two-compartment environment containing nutrients permitting growth in one region followed by an antibiotic region [7, 29]. We study the decay of the population density in the antibiotic region (as shown in 3 a). In that region, the populaiton dynamics is described by Equation (2) with growth terms replaced by death terms (with negative growth rates 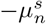 and 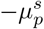 for the normal and persister subpopulation, respectively). We consider the extent to which the wave spreads into the unfavorable environment by calculating the penetration distance *x_e_* into the antibiotic compartment. Numerically, it is computed as the distance of the position where *n*(*x_e_*) + *p*(*x_e_*) = *N*_tip_ (*N*_tip_ = 0.05 is used for the simulations). Simulation results are shown in 3 b). The penetration depth of the wave depends on the phenotype switching rates and displays a maximum for intermediate phenotype switching rates (3 b). In the absence of phenotype switching, the normal population decays exponentially (as 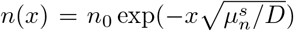) in the antibiotic region and the penetration depth is given by 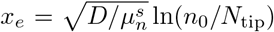. For finite switching rates, the penetration depth increases due to an increase in the fraction of persister cells at the boundary, which survive longer and travel over larger distances when they enter the antibiotic region. If the switching rates are equal (*a = b*) as in 3 b), for fast phenotype switching rates, persisters switch back to the normal cell state inside the antibiotic region, which causes effectively an increase in their death rate and thus a decrease in the penetration depth. For small phenotype switching rates, the extent of the wave is initially small but increases as the phenotypic redistribution occur at longer times.

These considerations can be made more precise by analyzing the steady-state density profile in the antibiotic region. In the steady-state, the wave equations 2 result in diffusion-decay equations coupled through phenotype switching terms, which can be analytically solved (see the appendix). The analytical solution shows that the subpopulation decays exponentially with two characteristic decay constants *κ*_±_. The slow decay term with decay constant *κ*_-_ dominates far into the antibiotic region, where the total population profile can be approximated as

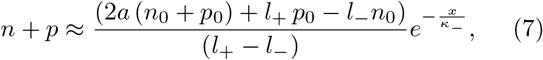

where *n*_0_ and *p*_0_ are the populations at the compartment boundary and *l±* and the decay constant *κ_-_* are func-tions of the growth, mobility and switching parameters as given in the Appendix (for *D_n_* = *D_p_* = *D*). From this expression of the profile, we obtain the penetration depth as

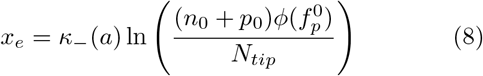

where 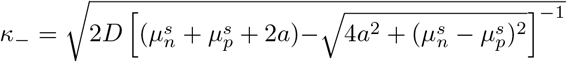
and

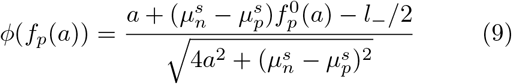

is a function of the parameters of the dynamics and of the persister fraction at the compartment boundary, 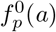. Therefore, the decay of the population wave in the antibiotic region is governed by two factors – the spatial decay constant *κ*_-_(a) and a function of persister fraction 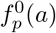 at the interface between growth and antibiotic, both of which are a function of phenotype switching rate. The decay constant *κ*_-_ is a decreasing function in *a* (as shown in Figure 4b) whereas *f_p_*(*a*) is an increasing function (as shown in Figure 4a). The combination of these two opposing behaviors explains the non-trivial maximum in the extent of the wave with the variation in phenotype switching rates. In Figure 4c, we compare the analytical expression of the extent of the wave to the numerically computed value.

**FIG. 3:**
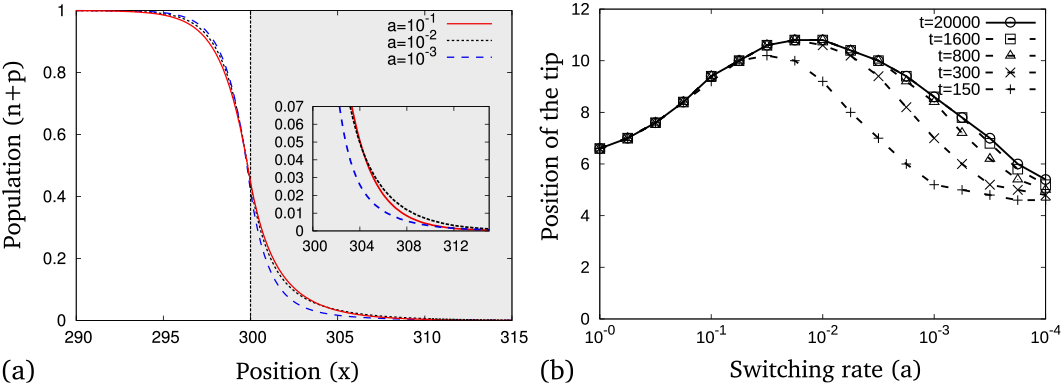
(a) Steady-state profile of the total population inside an antibiotic region (grey area) for different phenotype switching rates, *a* = *b* = 10^−1^ (solid red line), 10^−2^ (black broken line), 10^−3^ (dashed blue line). The inset shows that the population decay is the slowest for an intermediate phenotype switching rate. (b) The penetration depth of the population wave into the antibiotic region at different times as a function of the phenotype switching. The plot shows the existence of a phenotype switching rate for which the extent of the wave into the antibiotic region is maximal.

**FIG. 4:**
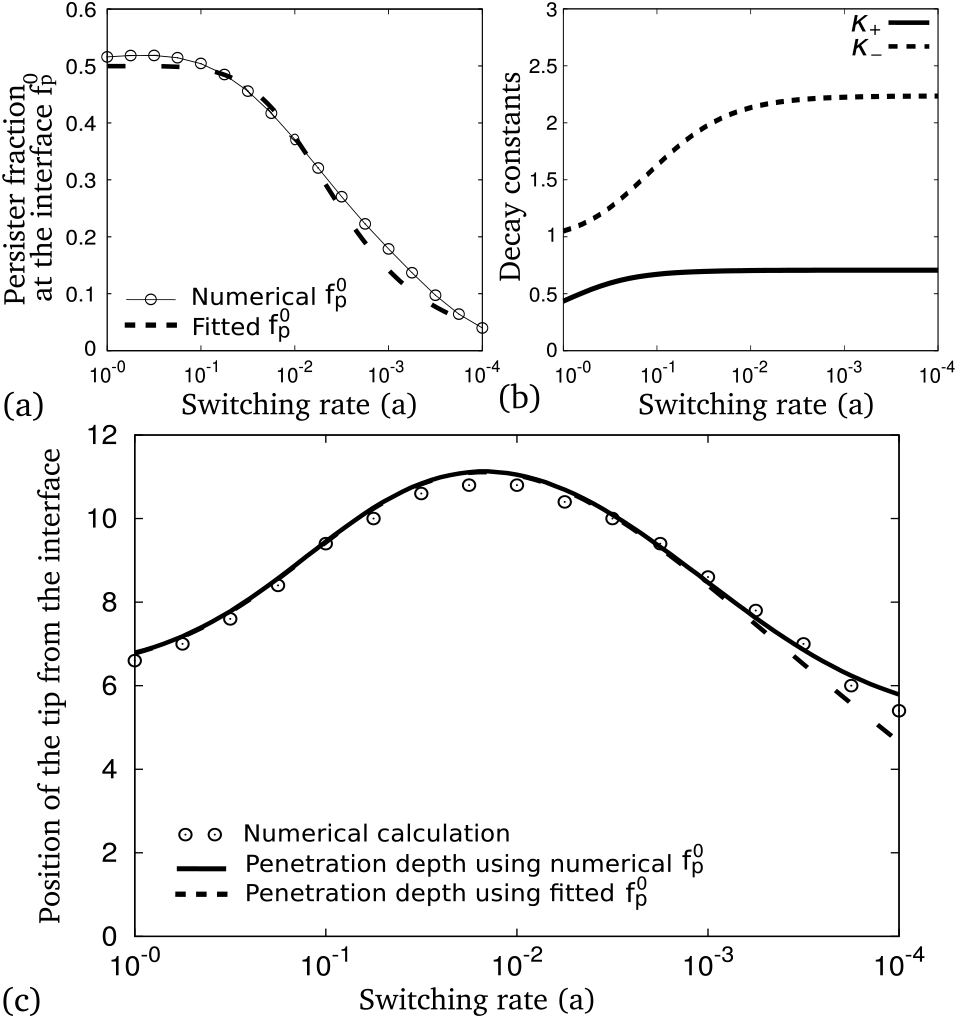
(a) Persister fraction at the boundary of growth and antibiotic regions as function of the phenotype switching rate (open circles). For large switching rate, the persister fraction approaches the maximal value i.e., *f_p,max_* = *a*/(*a* + *b*) (= 0.5 for equal switching rates). The dashed line shows a function that approximates the variation of fp with a. The fit function is 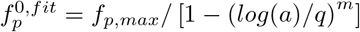 with q=2.5 and m=5. (b) Two characteristic decay constants of the population density in the antibiotic region, *κ*_+_ (solid line) and *κ* (dashed line). Both are increasing functions of the switching rate. (c) The impact of the switching rate on the persister fraction at the boundary and on *κ* are sufficient to explain the maximum in the extent of the wave as a function of the phenotype switching rate (open circles are from the numerical calculation). The solid line uses Eq. 8 together with the numerical values of 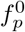, whereas the dashed line uses the fit function from (a).

### Crossing of an antibiotic barrier

The extended distance over which the population decays within the antibiotic region due to the presence of persisters can also support the crossing of such a region of finite width, i.e. an antibiotic barrier, a scenario we have previously studied in a model with discrete spatial compartments [21]. We found that the presence of persister cells can decrease the mean first arrival time of cells in a growth environment behind an antibiotic barrier. The same is seen in the continuous-space model described by the two-subpopulation Fisher equation that we study here. To study this scenario, we numerically calculate the crossing time of expanding population facing a spatially-extended antibiotic barrier of finite width (the schematic is depicted in 5 inset, we determine the crossing time as the time between the arrival of the tip of the wave at the first and second interface). We find the crossing time to show a non-monotonic behavior as a function of the increase in phenotype switching rate, as shown in Fig. 5. Specifically, we observe a minimum at intermediate switching rates that reflects the maximum of the penetration depth into the antibiotic region discussed above. At very low switching rates, as one would expect for a population with no persisters, the crossing time scales with the width of the barrier and their ratio approaches similar values as shown in Fig. 5. Similarly for very fast switching rates, the persister population first crosses the barrier, and hence the crossing time again scales with barrier width. For intermediate switching rates, the ratio of crossing time and barrier width depends both on the rate of phenotype switching and the width. This is because both normal and persister subpopulation contribute to barrier crossing but at different length scales; the normal cells dominate crossing over narrow barriers, while the persister cells enable crossing of wider barriers.

**FIG. 5:**
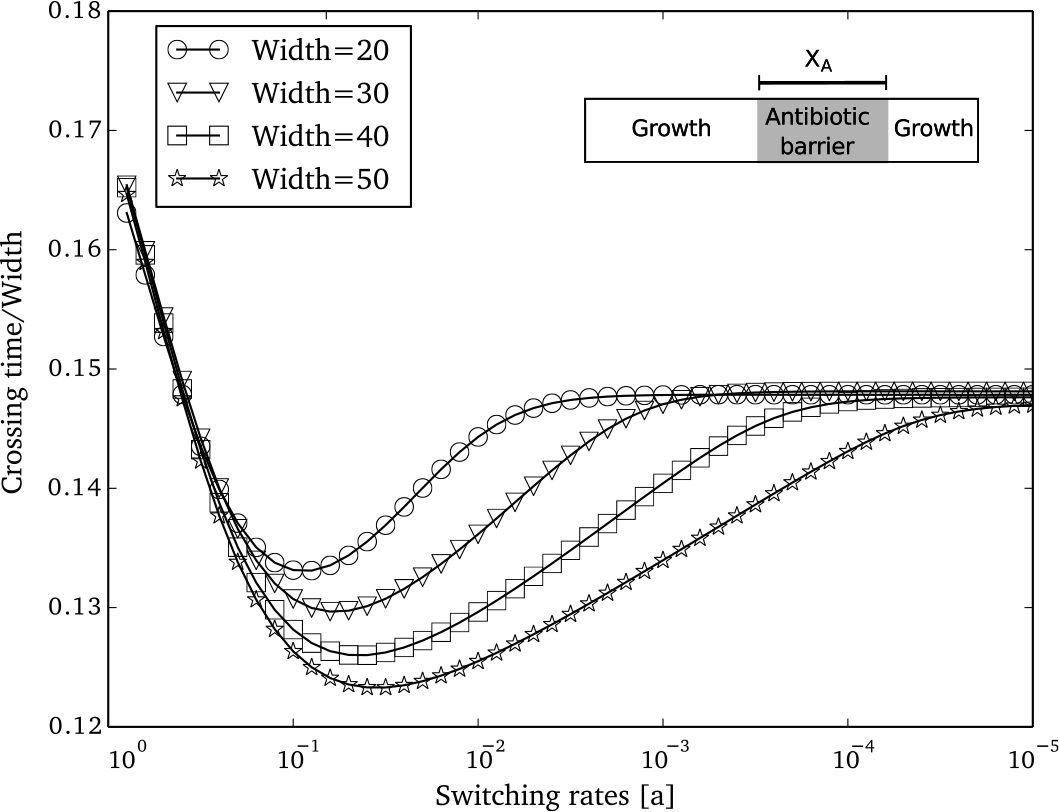
Antibiotic barrier crossing time of the population as a function of the phenotype switching rate *a* for different barrier widths. The crossing time is normalised with respect to the barrier width.The inset schematic depicts the simulation setup. A long growth region is chosen to allow the population to reach a steady state (for each *a* value) before facing the antibiotic barrier.

## DISCUSSION

Theoretical and experimental studies have shown stochastic switching between distinct phenotypes in bacterial population as a bet-hedging strategy for temporally varying environments [18–20, 30–33]. A recent study showed bet-hedging is more favorable in spatially varying environments compared to temporally varying environments [34]. However it is unclear how stochastic switching dictates bacterial growth in spatially varying environment. Here we have used a Fisher wave approach to investigate the spatial expansion of a bacterial population with stochastic phenotype switching for two scenarios, a spatially homogeneous environment and an antibiotic barrier. The cost of the presence of persisters during growth-favorable conditions is quantified by the population expansion speed, for which we obtained analytical expression. For typical switching rates, this cost is very small, even if there is a substantial persister fraction in behind the wave front, because the tip of the population wave contains a rather small fraction of persister cells. The persister pool in the back of the wave acts as a reservoir for the case of encountering stressful environments. The subpopulation redistribution from the tip to the back of the wave occurs at a slower rate than the population expansion (discussed in the Appendix). This contributes to the low cost associated with persistence. At an boundary to a stress environment such as an antibiotic barrier, the sub-population structure has to catch up with the wave tip, which results in transient stalling of the expansion. Eventually, the number of persister cells at the boundary increases due to subpopulation redistribution, helping the population wave to spread into the antibiotic region, thus providing the fitness advantage conferred by the presence of persisters. Moreover, we find an optimal phenotype switching rate for most rapid barrier crossing. The maximum occurs due to the subpopulation distribution at the barrier interface and the switching of persister cells to the normal phenotype. The existence of optimal switching rates for barrier crossing is reminiscent of optimal switching rates for growth in a temporally varying environment [18–20] In summary, our study reveals an added advantage of bacterial persistence in spatial environments which may play an important role in bacterial invasion and in the development of antibiotic resistance, where persisters may provide a pool from which resistance can emerge.

## APPENDIX

### Phenotypic redistribution in interior of the expanding wave

To quantify the change in phenotypic redistribution from the leading edge to the interior of the wave, we define a phenotypic flux as 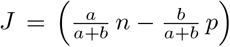. The phenotypic flux balances out in the interior of the wave, i.e., *J* = 0. In the moving frame of reference (*z = x* — *ct*), the phenotypic flux increases (and persister fraction decreases) from zero at the interior of the wave to the front of the wave. The increase in J inside the fully populated region (where *n + p* = 1) is governed by a diffusion-decay equation,

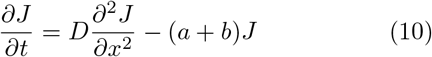

For an expanding population, the phenotypic flux in the moving frame of reference thus increases as

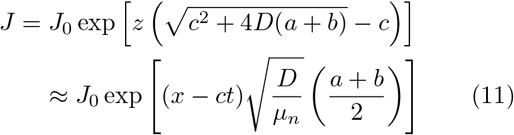

The above expression shows that subpopulation redistribution occurs at a slower rate (with rate constant 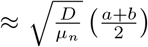 for small *ab*) and that the subpopulation redistribution process lags behind the advancing wave as shown in Figure S1. Therefore for small switching rates, spatial expansion results in keeping fraction of persisters at the tip of the wave small despite a large fraction of persisters in the interior of the wave.

### Wave speed determination through traveling wave analysis

Using the wave solution ansatz

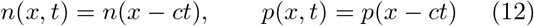

and introducing auxiliary variables *n*’ = *dn/dz* and *p*’ = *dp/dz* with *z = x* — *ct*, equations 2 can be expressed as the following set of autonomous first order equations,

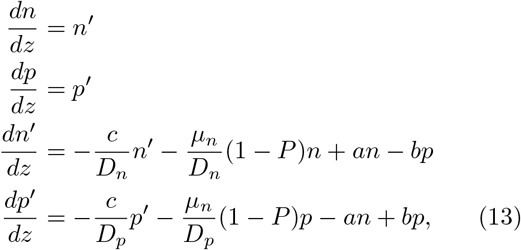

which can be analyzed by the standard fixed point analysis for traveling wave solutions. The eigenvalues of the

**FIG. S1:**
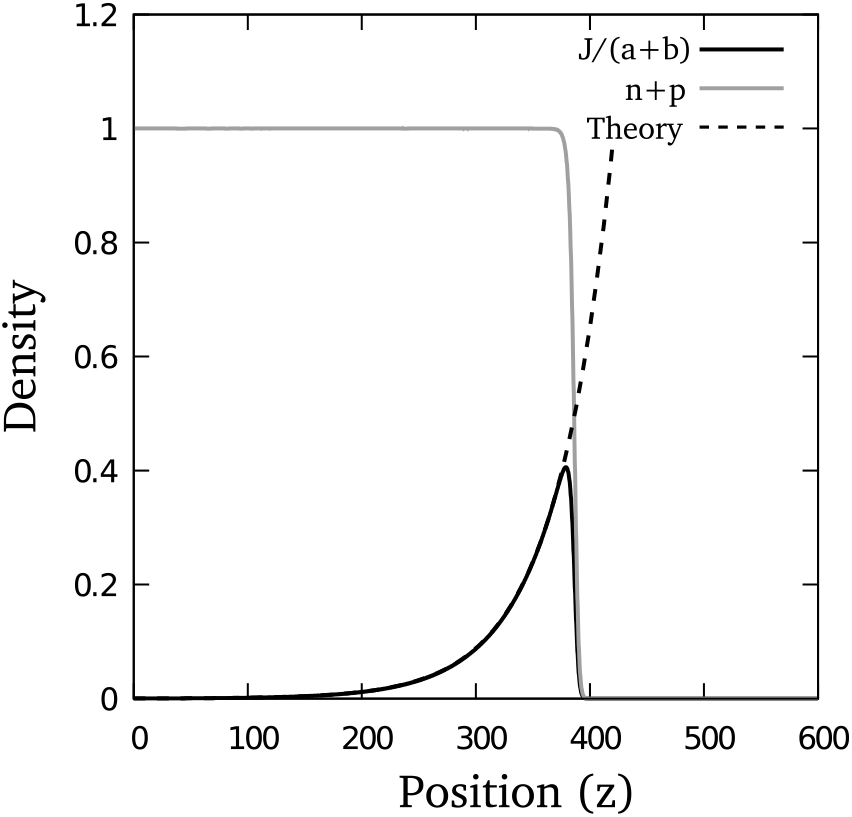
Phenotypic flux distribution along the wave in the moving frame of reference (*z*). In the populated regions (where *n* + *p* = 1), the phenotypic flux *J* (solid black line) increase exponentially from the back of the wave to the front, in excellent agreement with the analytical expression for *J* (dashed black line).

above equations near the fixed point *n*’ = 0,*p*’ = 0,*n* = 0,*p* = 0 is given by the following fourth-order equation

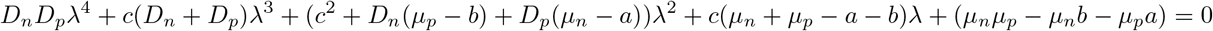

For simplicity, we take *D_n_* = *D_p_* = *D*. The four roots(λ_1,2,3,4_) are then given by

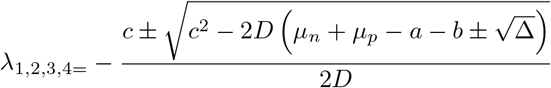

with Δ = (*a*+*b*)^2^ + (*μ_n_* — *μ_p_*)^2^ — 2(*μ_n_* — *μ_p_*)(*a — b*). For the existence of a stable traveling wave solution, these these eigenvalues must be real. This leads to the condition,

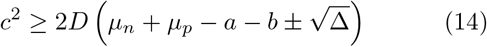

which for *a* = *b* simplifies to

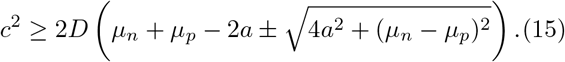

In the limit *a* = *b* = 0, the two conditions result in the marginal speeds of two uncoupled waves, 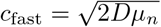 and 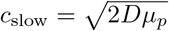. In the coupled case, the wave speed is given by the faster moving wave, i.e. the positive root.

### Population decay in the antibiotic region

In the antibiotic region, the steady state equations simplify to two coupled diffusion-decay equations,

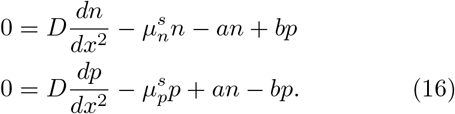

These equation can be solved for boundary conditions *n*(*x_b_*, 0) = no and *p*(*x_b_*, 0) = *p*_0_, from which we obtain the following subpopulation profile (for *a* = *b, D_n_* = *D_p_* = D):

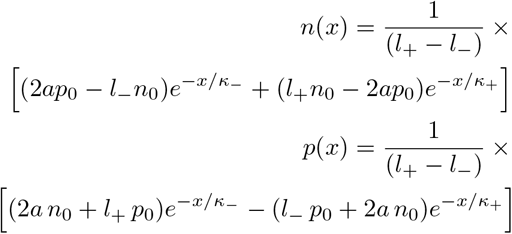

with 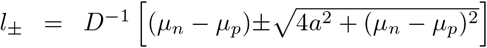 and 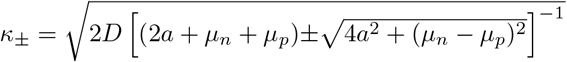.

